# Decoding Individual Musical Pitches in Perception and Imagery using Evoked Theta and Beta Power in EEG

**DOI:** 10.64898/2026.07.19.736806

**Authors:** Mi-young Chung, Charlotte Van Barr, Andrea R. Halpern, Robert J. Zatorre

**Author notes:** **Correspondence to:** Robert J. Zatorre, Montreal Neurological Institute, McGill University, Quebec, NW230, 3801 rue University, Montreal, Quebec, H3A 2B4.

## Abstract

Imagery evokes perceptual-like experiences and recruits similar neural activity to perception. In music, this shared activity has already been established in secondary auditory cortex, yet the nature of the representation is not fully understood; specifically, it is not clear whether the identity of individual imagined pitches can be decoded from brain activity. Using temporal lobe EEG channels, we examined the shared evoked patterns of musical pitch perception and imagery, and decoded individual imagined pitches from evoked oscillatory power across three conditions (Perception, Imagery, Random Pitch). Twenty-two participants heard the opening tones of a familiar melody, imagined its continuation, and judged whether a final probe tone matched the original note; a random-pitch condition served as a control. Replicating previous findings, the evoked perceptual pattern was reinstated during imagery, particularly in the right temporal channels, which showed stronger event-related potential correlations and a larger mismatch negativity to the probe tone than the left. A multiclass support vector machine decoded individual pitch identity from evoked oscillatory power (phase-locked activity time-locked to each tone onset) exceeding chance in all conditions, with accuracy higher for Perception and Imagery than Random Pitch. The optimal oscillatory bands differed across conditions: theta-beta gave the highest accuracy for Perception and Imagery, theta-alpha for the Random Pitch. These results demonstrate that individual pitch information is present during both perception and imagery, and that the theta–beta signature shared by the predictive contexts, distinct from the control, suggests a contribution of predictive processing.

**Significance Statement:** Musical imagery — the experience of “hearing” music in the mind’s ear — recruits brain activity resembling perception, yet whether the pitch specific information during imagery can be read out from this activity has remained unknown. Using EEG, we show for the first time that individual imagined pitches can be decoded from evoked oscillatory power, and that perception and imagery share a spectral signature (theta–beta) absent from a non-predictive random-pitch control. This dissociation indicates that imagined pitch reflects top-down predictions reactivating sensory representations, consistent with a predictive-coding view of imagery. These findings clarify the oscillatory basis of musical imagery and establish that its specific content is recoverable from non-invasive signals, informing future imagery-based brain–computer interfaces.

## Introduction

Imagery is a fundamental human cognitive process that enables replay of internal representations of sensory experiences, raising critical research questions about the nature of the information represented. Music provides an interesting test case for the study of imagery because of its multifaceted nature, including pitch, temporal, and timbral information. Although we know that representation of pitch information is critical for many aspects of music cognition (Peretz, 2016; Tillman et al., 2023), it is not clear whether or how specific tonal pitch information is encoded in neural activity during imagery.

Studying auditory imagery remains challenging due to its covert and temporally dynamic nature, which makes it harder to design stable behavioral markers compared to visual imagery (Hubbard, 2018; Cohen et al. 2009). However, converging evidence from diverse experimental approaches, including functional imaging and lesion studies, shows that auditory perception and imagery similarly depend on the secondary auditory cortex, particularly in the right hemisphere (Zatorre and Halpern, 2005). More recent fMRI work shows that time-resolved activity patterns associated with listening to melodies are reinstated during imagery within auditory cortex (Regev et al., 2021), indicating that neural representations during imagery contain information about specific melodies. Furthermore, using MEG Herholz et al. (2008) showed that the mismatch negativity (MMN) evoked by imagery resembles the perceptual MMN, indicating that predictive processes are involved in both perception and imagery. That study also showed that musicians exhibited stronger MMN responses in the right auditory cortex during imagery, in keeping with other perceptual MMN studies (Tervaniemi et al., 1999; Bonetti et al., 2022).

To delve more specifically into the nature of imagery for pitch information, prior studies have examined temporal and oscillatory electrical activity in the brain. Janata (2001) reported comparable N1 responses during imagined and perceived tones using EEG, with decreasing amplitude patterns across successive imagined pitches within a trial, suggesting auditory top-down processing (Navarro and Janata, 2010; Näätänen et al., 2007). Oscillatory analyses have further highlighted the role of low-frequency bands (<30 Hz) which are associated with top-down processing (Keil and Senkowski, 2018). Gelding et al. (2019) reported increased beta-band activity in sensorimotor areas and theta-band interactions between somatosensory and auditory areas that support accurate pitch imagery. These findings suggest that low-frequency oscillations in the auditory cortex are crucial for musical pitch imagery, although how neural responses represent pitch during imagery remain unclear.

Decoding analyses have recently advanced imagery research further, by applying inference of hidden states to understanding covert brain activity (Wolff et al., 2017; Loriette et al, 2022). In musical imagery, decoding studies have captured internal representations of scale degrees (a note’s function within a key) and melodic structures in the auditory cortex using fMRI and MEG (May et al., 2023; Quiroga-Martinez et al., 2024). However, few studies have directly examined decoding of pitch information (absolute frequency of the sound) during imagery. Chung et al. (2023) used EEG and demonstrated successful decoding of seven separate pitches during imagery, introducing the potential to decode specific pitch information, although the task design did not permit separation of visual cues from imagery.

The present study investigates how individual pitches are internally represented in the brain during a musical imagery task. Using seven pitches across three familiar melodies, we examined EEG activity in musicians without absolute pitch, employing a paradigm adapted from the Herholz et al. (2008) study. EEG was chosen for its high temporal resolution, allowing for precise decoding across time and frequency domains, with temporal-lobe channels selected based on auditory evoked potential amplitude We hypothesized that: 1) imagery and perception share similarities in evoked potential patterns and MMN responses in auditory cortex, particularly in the right hemisphere; and 2) individual pitch values during imagery are encoded in low-frequency oscillations (Gelding et al., 2019). To test these hypotheses, we employed both univariate analyses, including ERP and MMN, alongside multivariate decoding using SVM classifiers trained on various combinations of oscillatory frequency from evoked activity.

## Materials and Methods

### Participants

Twenty-eight musicians were initially recruited for this study. They were selected based on a set of inclusion criteria: (1) musicians with a minimum of five years of formal music training, because individuals with musical expertise tend to have more refined auditory imagery skills, particularly for pitch, which contributes to higher signal-to-noise ratios in neural measures of auditory processing (Herholz et al, 2008); (2) not having Absolute Pitch (AP) due to the distinct neurological representations of pitch information in individuals with AP compared to the general population (Burkhard et al., 2020); (3) familiarity with the three melodies used in the experimental task; (4) normal hearing, good overall health, and aged between 18 and 40 years; (5) passing the pre-test consisting of the behavioral task of the experiment (See Behavioral Task and the Procedure, below). Their musical backgrounds and auditory imagery abilities were quantified via two validated questionnaires, Montreal Musical History Questionnaire (MMHQ) (Coffey et al., 2011) and Bucknell Auditory Imagery Scale (BAIS) (Halpern, 2015). Five participants were screened out based on the pre-test and an additional one was excluded in the data analysis due to poor signal quality. The final cohort of 22 subjects (8 male, 14 female, mean age 24.73±4.7) involved in this study had an average of 13.59 ± 5.72 years of professional music training, ensuring a high level of expertise in auditory processing. Average BAIS Vividness score was 5.714 ± 0.746 and BAIS Control was 6.013 ± 0.746 (In total 5.864 ± 0.672), exceeding the standard average of 5.1 for musicians (Halpern, 2015). The research protocol was approved by the Research Ethics committee of the Faculty of Medicine and Health Sciences at McGill University.

### Stimuli

Three familiar melodies (Figure 1) were selected for the study, on the basis of their pitch composition (Table 1), tonality, and rhythm covering an appropriate range for EEG-base decoding. The melodies were altered slightly to ensure that they contained isochronous pitches separated by silences of the same durations as the tones. The pitch composition was determined to guarantee a sufficient number of trials were presented for imagery decoding: more than 100 trials for each pitch in imagery. However, two tones, B3 and C3 were not sampled during the perception task; to compensate for this issue, a separate control task was presented (see below). The individual pure tones of the melodies were sinusoids in the range of 130.81 to 246.94 HZ, covering the diatonic musical scale from C3 to B3, and were created using Reaper (Cockos, Inc., USA). The duration of each tone was 400 ms and a fade-in/out of 300 ms was applied to avoid transients by synthesizing the sound signal in MATLAB. The inter-tone interval was 400 ms.

**Figure 1.**
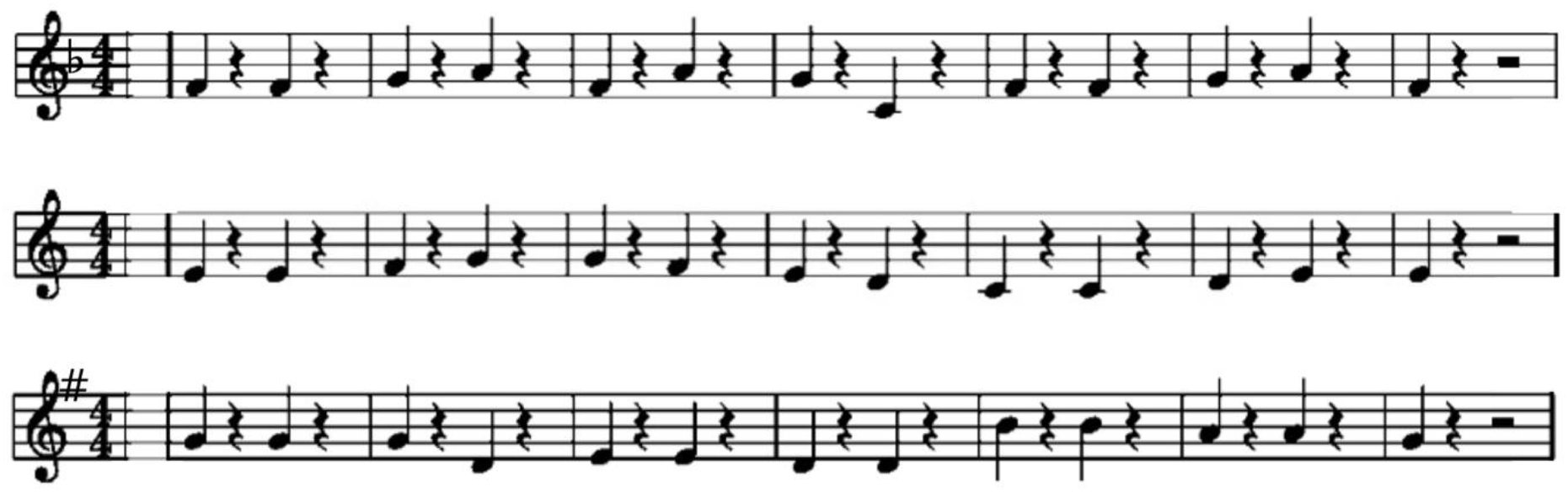
Three Familiar Melodies: Yankee Doodle (top), Ode to Joy (middle), and Old Macdonald had a Farm (bottom) in musical notation. Tones used in the experiment were presented one octave below those notated here.

**Table.** Number of times each of the seven tones of the diatonic scale appears for each condition using three melodies. First two columns describe the note distribution after one trial summed across melodies; the other two columns describe the total number of each tone across the entire test session (50 trials per each melody).

| Diatonic scale<br>tone name | Perception | Imagery | Perception_total | Imagery_total |
| --- | --- | --- | --- | --- |
| C3 | 0 | 3 | 0 | 150 |
| D3 | 1 | 4 | 50 | 200 |
| E3 | 4 | 2 | 200 | 100 |
| F3 | 5 | 2 | 250 | 100 |
| G3 | 6 | 2 | 300 | 100 |
| A3 | 2 | 3 | 100 | 150 |
| B3 | 0 | 2 | 0 | 100 |

To facilitate synchronization of auditory imagery with each pitch of the melodies, we presented a white dot flashing at the center of the screen synchronized with each tone throughout the task (Figure 2), allowing for phased-locked EEG analysis. The dot was synchronized with the tone onset and offset (400 ms, with 400 ms inter-stimulus interval), and was identical for every tone, conveying no information about the pitch value.

**Figure 2.**
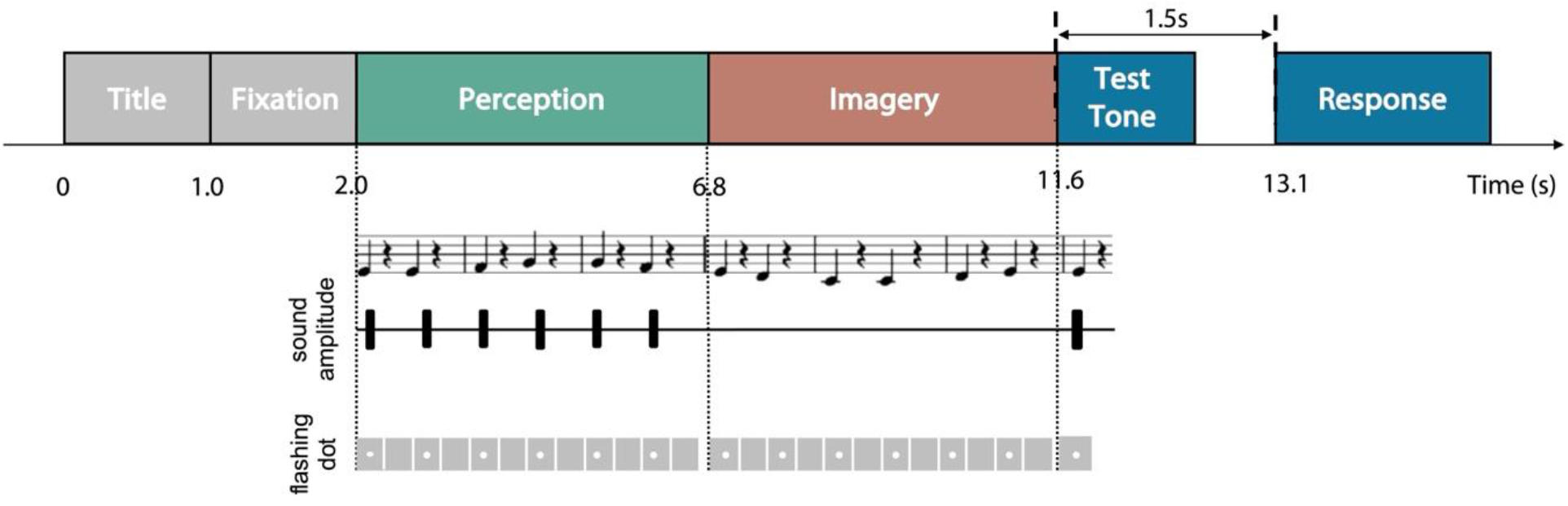
Behavioral task. One trial of the behavioral task consists of presentation of the song title, fixation point, perception phase (during which tones were sounded), imagery phase (during which no tones were sounded), and tone-matching (test-tone) phase. The flashing white dot was presented during the perception and imagery phases, in synchrony with the sounded or imagined tone.

### Behavioral Task

Each trial of the behavioral task involved the presentation of the title of the melody (e.g. “Yankee Doodle”) for 1 s, a fixation cross for 1 s, and the melody task for around 14 s (Figure 2). The melody task was adapted from Herholz et al. (2008), and each trial had three phases: Perception, Imagery, and Tone-matching (test tone). During the Perception phase, the participant listened to the first segment of the melody, consisting of six notes, while also viewing the synchronized flashing dot. In the Imagery phase no sound was presented but the participants were instructed to imagine the continuation of the melody – comprising the next six notes – while synchronizing their mental auditory representation with the timing of the flashing dot. The final phase, Tone-matching, involved the presentation of a single test tone, which either matched the last tone of the melody or differed from it one step or half-step in pitch (higher or lower). Participants were asked to indicate whether the test tone matched the last note of the melody or not by pressing a key on the keyboard. This behavioral task was administered during both the pretest, during the screening process, and as the main test in the EEG recording session.

### Control (Passive) tasks

In addition to the main active task, two control tasks were administered. The first control task involved the presentation of the visual stimulus displayed on the monitor without any accompanying sound or imagery cues for 2 minutes (150 trials). to take account of any activity due solely to the visual input. The second control task (termed Random Perception) involved passive presentation of all seven tones from the main task but in random order and without any task instruction or behavior, but accompanied by the visual stimulus as in the main task. This task provides a baseline to compare against anticipatory processes that likely occur in the context of well-known melodies. Each tone was presented 50 times in a random order, 350 trials in total.

### Procedure

Participants were provided with detailed information about the experiment, including an overview of the experimental process and the behavioral task, and were given sufficient time to familiarize themselves with the three melodies and to practice the behavioral task to ensure they were comfortable with the procedure before the pre-test began.

The pre-test serving as the screening consisted of 12 trials of the behavioral task, with each melody presented four times in random order. To proceed to the EEG recording session, an accuracy rate of at least 80% was required. If this accuracy threshold was not met the first time, additional pretests (up to three) were given, and participants qualified if they showed improvement in performance across trials.

Following the pre-test, the participants proceeded to the EEG recording session. The session began with the presentation of the two passive tasks. After completing the passive tasks, the participants performed the main test session. This main test consisted of 150 trials (50 trials per melody) of the behavioral task, and 75 trials were for ‘Match’, another 75 trials were for ‘Mismatch’ with test-tone of one step or half-step higher or lower. The experiment was divided into 5 blocks (30 trials per block) and short breaks of up to 2 min between blocks were given for rest and to maintain focus.

### EEG Data Recording

EEG signals were recorded using an EEG amplifier (BRAINAMP MR PLUS, Brain Vision, GmbH, Gilching, Germany) and 64 active electrodes (actiCAP, Brain Vision, GmbH, Gilching, Germany). The 10-10 system of the American Clinical Neurophysiology Society guideline 2 montage was utilized with the reference electrode on the right cheek (Acharya et al., 2016). The impedance of all 64 active electrodes was verified to be below 5kΩ to ensure optimal signal quality.

### Data Analysis

The EEG data analysis was carried out using the EEGLAB toolbox implemented in MATLAB. First, the EEG data were resampled to 200 Hz for more efficient computation. Then, the standard pipeline for EEG preprocessing was applied, as follows (Bigdely-Shamlo et al., 2015). The band-pass filter was applied with a passband of [1, 50] Hz. Bad channels were identified and rejected using kurtosis and interpolated using the spherical algorithm. The data were re-referenced to both mastoids, chosen based on their verified effectiveness in auditory-related experiments such as MMN studies (Mahajan et al, 2017), then the REST algorithm was applied to standardize the reference channel at the mastoids to infinity (Dong et al., 2017). Independent Component Analysis (ICA) was employed to remove the artifact using the Second-Order Blind Identification (SOBI) algorithm, as it has been shown to improve artifact removal while maintaining computational efficiency and accuracy, particularly in removing ocular and other non-neural artifacts among the variety of ICA algorithms (Urigüen et al., 2015).

After the standard EEG preprocessing aiming artifact removal, ICA was re-applied to eliminate potential visual-related interference from the flashing dot stimuli. This was accomplished using data from the visual control session. The procedure involved the following steps: 1) Independent components were extracted from the visual control session data using the SOBI algorithm; 2) Correlations were calculated between the power spectrum of these independent components’ activities and the brain activity recorded from temporal channels during the first block of the task; 3) Independent components were selected for removal if their correlation exceeded a threshold defined as [((mean of the correlation values)+0.75)*(standard deviation of the correlation values)]; and 4) the spatial filter, derived by combining Blind Source Separation and Regression a method recognized for its efficacy in artifact removal in EEG analysis – was applied to the data (Guarnieri et al., 2018). This approach allowed for the attenuation of overlapping brain activity between the visual control session and the main task. The efficacy of this method in attenuating visual components is illustrated in the supplementary materials (Figure S1).

Only trials with correct responses during the tone-matching phase in the behavioral task were included in the data analysis. Including only accurate responses is essential, as that provides evidence of time-locked musical pitch imagery synchronized with each tone onset, ensuring the phase-locked EEG activity to individual pitch imagery.

The analysis of the EEG data primarily focused on Left and Right temporal channels (Figure 3). The validity of the channel selection for auditory cortex was verified via inspection of the expected N1 activity to the sounded tones (see Results). After analyzing the temporal lobe channels, additional channels of whole brain EEG electrodes and all channels within each hemisphere were also considered for the decoding analyses, to see if they contained additional useful information beyond those from the temporal lobe channels.

**Figure 3.**
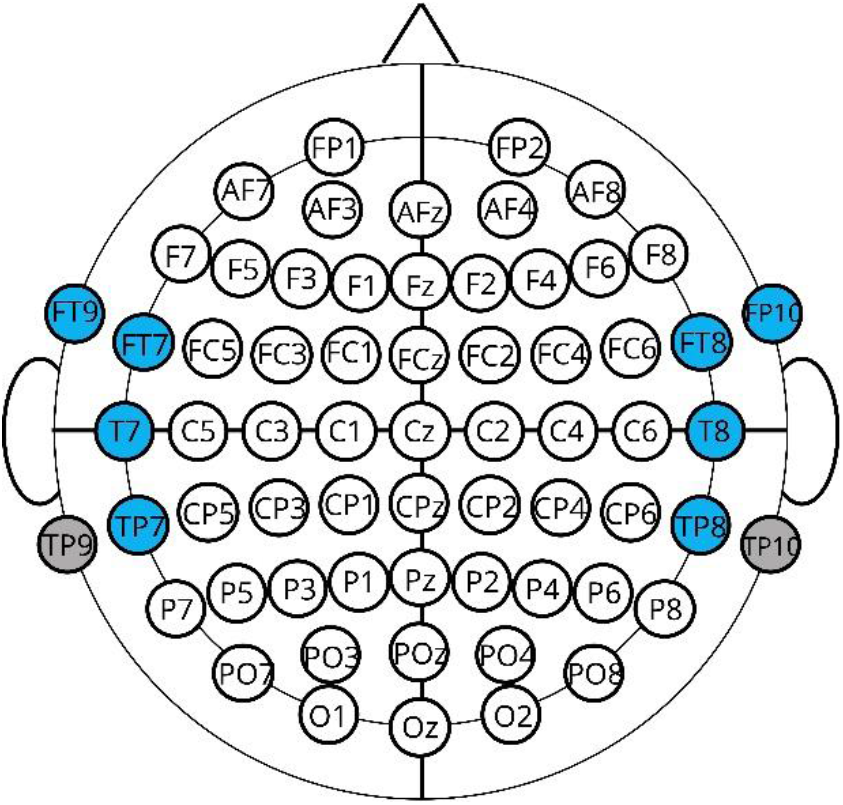
EEG channel montage. The channels for the temporal lobe are colored blue: for left hemisphere FT9, FT7, T7, and TP7, and for right hemisphere FT10, FT8, T8, TP8. Channels with grey color are the mastoids channels used as the re-reference channels in the preprocessing.

Event-Related Potentials (ERPs) were computed by applying a low-pass filter of 13 Hz and by segmenting into epochs from −300 ms to 500 ms relative to stimulus onset. Each epoch was baseline corrected with the baseline from −300 ms to 0 ms, z-score normalized, and then categorized according to the tone and task. Note that only the data during the behavioral task were used here, and only ERPs for 5 pitches (D3 to A3) were employed to compare Perception and Imagery. To confirm the evoked potential during imagery, we compared the ERP amplitude after the stimulus onset to the 95% confidence interval estimated from the baseline. The time range of this exceeding evoked potential was 110 ∼ 190 ms, which was selected as the time window for N1 corresponding to the previous literatures (Janata, 2001).

The imagery Mismatch Negativity quantifies the difference in neural activity between instances when the test tone matches the melody and when it does not (Herholz et al., 2008), which validates the auditory imagery of participants. To calculate imagery MMN, the ERPs corresponding to match trials were subtracted from the ERP of the mismatch trials. The confidence level (95%) estimated from the baseline of imagery MMN was applied to determine the variance of imagery MMN distribution over time.

### Evoked Power Analysis and Multiclass Support Vector Machine (Multi-SVM)

The evoked power is the phase-locked spectral activity to the stimulus onset. We used the epoch segmented from −300ms to 500ms relative to stimulus onset, for the evoked power analysis. To calculate the evoked spectral power, the total spectral power was computed and subtracted from the induced power. This induced power is computed by calculating the spectral power from the epochs that the ERP was subtracted from. Both total power and induced power were calculated by Short Time Fourier Transform with a 100 ms window size with 90% overlap. For the evoked power for each pitch during Perception, Imagery, and the Random Perception task, the frequency band power was computed by averaging across four frequency bands: theta (4-8 Hz), alpha (9-12Hz), beta (13-30 Hz), and gamma (31-50 Hz).

Multiclass Support Vector Machine (Multi-SVM) was employed as the classifier model for the five-pitch classification in the Perception and the seven-pitch classification in the Imagery and the Random Perception conditions. The input data for the Multi-SVM were the evoked frequency power of band groups: theta-alpha pair; theta-beta pair: alpha-beta pair; lower frequency bands (theta-alpha-beta), and all bands(theta-alpha-beta-gamma), extracted from the temporal electrodes. The decoding analysis spanned the entire duration of each epoch, including the baseline period, by training and testing the model every 10 ms with a 100 ms duration sliding time window. To evaluate the performance in decoding pitches, the chance level of the decoder was estimated at each time point using permutation analysis, averaging the decoding accuracies across 100 iterations of a model trained on the same dataset but with randomized labels for the optimal band group for imagery. By using the same classification model configuration, we compared the results of band groups using the difference between the decoding accuracy and the chance level for each band group. Note that the chance level was computed with 10 iterations for each band group when comparing the band groups to avoid the excessive computation time. The significance of the decoding accuracy was compared with the chance level with confidence interval (95%) and cluster-based permutation test using baseline as a null distribution.

The Multi-SVM classifier was trained on 80% of the trials with Perception and Imagery data during the behavioral task, and the remaining 20% was used for testing. The trials were randomized for order then divided into training and testing sets. A 5-fold cross-validation procedure was applied for validation purposes.

Following the decoding of the temporal lobe electrodes, decoding results from the whole brain (all electrodes), and from left and right hemisphere electrodes separately were also analyzed to compare their decoding results with those corresponding to the temporal lobe channels, using theta & beta. This analysis assessed whether the classification accuracy of brain activity from both temporal lobes was similar or greater than from the whole brain, and whether there was a difference in accuracy for the left vs. the right hemisphere.

### Statistical Analysis

Statistical tests were utilized to verify: 1) the existence of an evoked potential and MMN during imagery as validation of the experimental design; 2) the predicted higher amplitude of ERP and MMN in the right temporal channels compared to the left, and 3) the expected similarity of spatial and temporal pattern during Perception and Imagery.

To verify the existence of N1 during Imagery, we compared the evoked potential after onset to the evoked potential during the baseline by comparing the evoked potential to a 95% confidence interval estimated from the baseline. This resulted in the decision of time window of N1. This confidence interval approach was also applied to the verification of MMN existence during imagery.

Then, a 2-way ANOVA test was employed with two factors: Hemisphere (Left vs Right) and Conditions (Perception vs Imagery vs Random Perception). For imagery MMN, statistical analysis was conducted on the peak amplitude using a mixed effect model with two fixed effects: Match/Mismatch and Left/Right hemisphere and a random effect: Subjects. This model was derived to determine if there was a significant difference between the match and mismatch condition, and between left and right hemisphere activities for all the correct trials, considering the imbalance of the correct trial numbers per participant. Also, paired t-test was used to determine the significance difference of MMN area under the confidence level (95%) between left and right temporal channels.

Pearson correlation analysis was utilized to test the similarity of evoked responses between Perception and Imagery. First, the correlation of the topographic distribution of grand averaged N1 between Perception and Imagery was calculated to test for any similarity of spatial patterns of evoked potential in Perception and Imagery. Then, we explored similarity between perception and imagery in the grand-average ERP responses with five pitches and compared the magnitude of the correlation between the left and the right hemisphere. Cross-correlation between perception and imagery with various time lags was also calculated for each hemisphere to measure any difference in time-locked brain activity in Perception and Imagery. ERP correlation analysis was held performed on the correlation between perception and imagery for subject with 2-way ANOVA test with two factors – 5 pitches (D/E/F/G/A) and Left/Right - after Fisher’s z-transform.

The comparison of decoding performance among band groups during each condition was carried out with a 2-way ANOVA test with 2 factors: Conditions and Band groups. The decoding performance was measured by the subtraction of the chance level from the actual decoding accuracy. Another 2-way ANOVA test was carried out for the lower frequency band pairs with 2 factors: Conditions and Time Period, to compare if there were difference in decoding performance among time period of Early (20∼120 ms), Middle (150∼250 ms), and Later (300-400 ms).

A 2-way ANOVA test was also employed with 2 factors: Conditions and Channel sets, to compare the performance of decoding models when different sets of channels were used (See Figure3): temporal channels, all channels in the left hemisphere (odd numbered channels), all channels in the right hemisphere (even numbered channels), and whole brain (all channels including midline channels).

## Results

Participants achieved an average accuracy level of 93.91±6.73% (range from 80% to 100%) in the behavioral task performance. This result validates that imagery was occurring and shows sufficient EEG data were available for a robust analysis.

ERPs were grand averaged to Perception and Imagery during the behavioral task, divided into left and right temporal channels (Figure 4). Average amplitude exceeding the 95% confidence interval estimated from the baseline showed significant negative potentials evoked from the stimulus, corresponding to the N1 ERP component, for both perception and imagery, exhibiting higher amplitude than the visual control session (Figure S2). The N1 during imagery showed a similar topographic pattern to perception (Figure 4), with the correlation of r = 0.82 (p = 2.48*10^−16^). Furthermore, the amplitude of the N1 was larger in the right hemisphere than the left (F(1,126) = 11.54 p = 9.09*10^−4^), with no significant difference among Conditions (F(2,126) = 2.24, p = 0.11), nor any interaction (F(2,126) = 0.18, p = 0.84, ANOVA).

**Figure 4.**
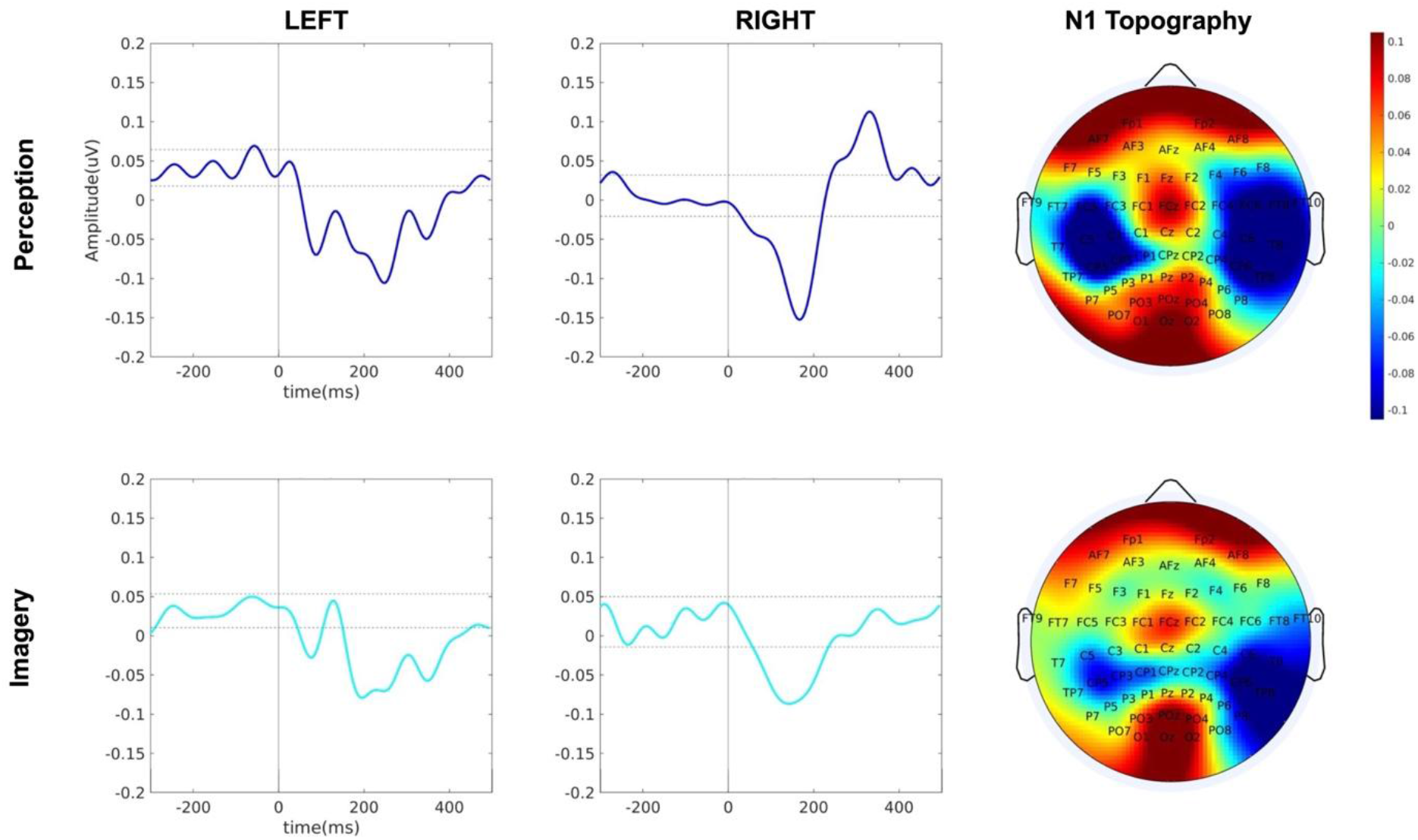
Grand average ERP patterns of Perception and Imagery in Left and Right hemisphere. 95% confidence interval estimated from the baseline for each condition is depicted with dot line. Topography of N1 component showing the bigger temporal channels’ activity. Above is for Perception and below is Imagery.

Similarity of ERPs for Perception and Imagery conditions was also demonstrated by the correlation analysis between the grand averaged ERP of perception and imagery in both left (r = 0.68 p = 8.01*10^−15^) and right temporal channels (r = 0.92 p = 3.17*10^−41^). The ERP time series for the two conditions were aligned with each other in time, as revealed by the cross-correlation analysis, which showed that the correlation between Perception and Imagery was highest when there was zero time lag both in left and right hemisphere (Figure S2). A 2-way ANOVA test using the Z-transformed values of correlation between Perception and Imagery of each participant revealed that the correlation was significantly higher in the right hemisphere than the left hemisphere (F(1,210) = 5.73, p = 0.018, η^2^_partial_ = 0.01), but there was no significant difference across pitch heights (F(4,210) = 2.01, p = 0.09, η^2^_partial_ = 0.02) nor was there an interaction (F(4,210) = 0.72, p = 0.58, η^2^_partial_ = 0.006).

Mismatch negativity from the test-tone ERP was observed both in the left and right hemisphere starting from 200 ms until 400 ms (Figure 5). The existence of a mismatch negativity and the difference between the hemispheres with the averaged amplitude during the time range of negativity revealed that the mismatch ERP was significantly more negative than the match ERP (F(1,6160)=4.29, p = 0.04, β = −0.09, mixed effect model), with higher amplitude in the right hemisphere (F(1,6160)=17.17, p = 3.46*10^−5^, β = 0.17, mixed effect model). The estimated effect was more negative in the right hemisphere vs. left hemisphere, but this effect did not reach statistical significance (F(1,6160)= 0.16, p = 0.69, β = −0.02, mixed effect model). However, the imagery MMN in the right hemisphere showed larger area under the 95% confidence interval estimated from the baseline when compared to the left hemisphere (Figure 5). This area under the 95% confidence interval was compared across participants, revealing bigger area in the right temporal channels (T(21) = 0.03, p = 0.01). Note that the time range for this area varied over individuals, averaged 325 – 370 ms for left hemisphere and 300 – 330 ms for the right hemisphere.

**Figure 5.**
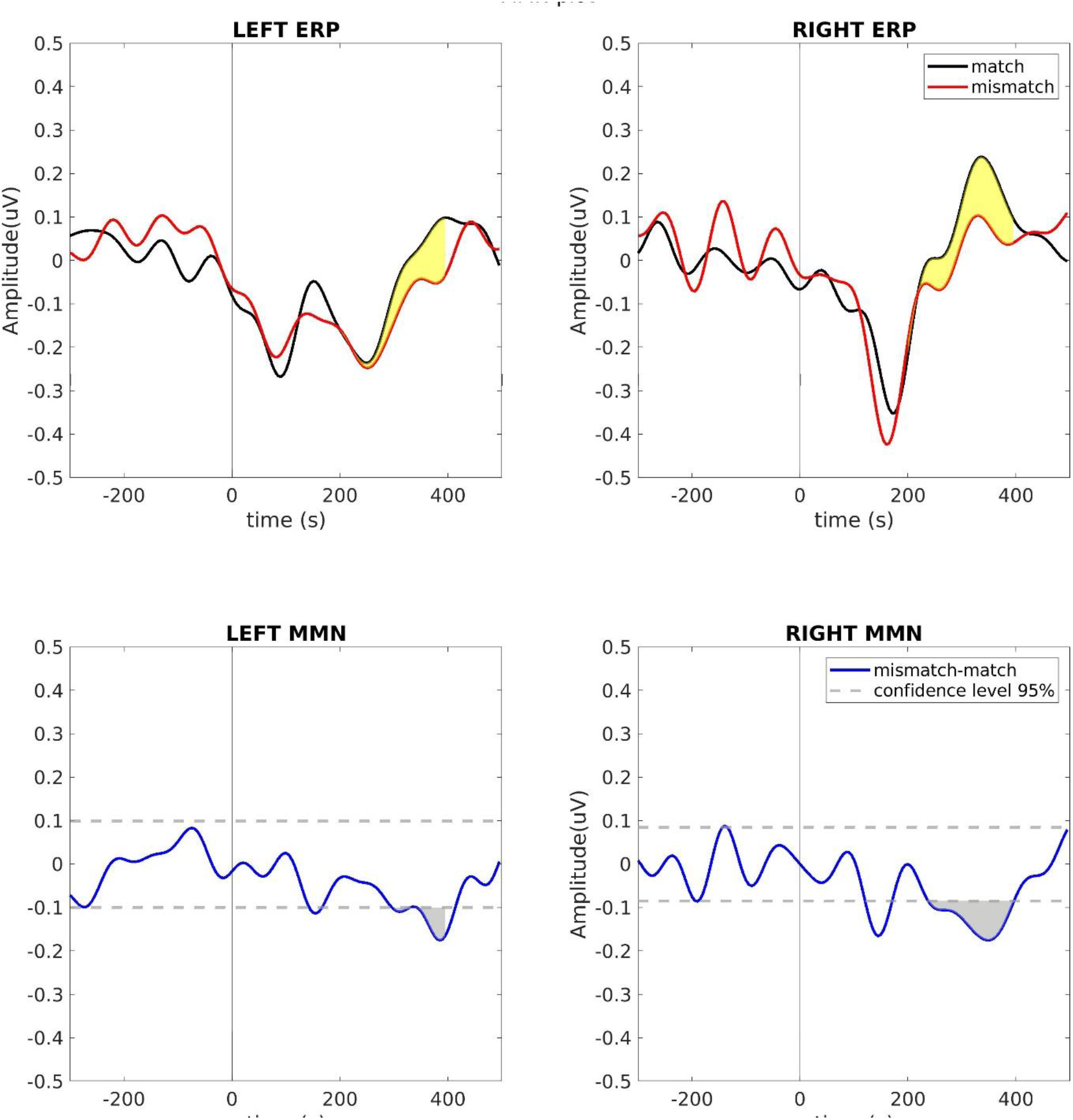
Imagery MMN. ERPs corresponding to match (black) and mismatch (red) are depicted in the upper row in temporal channels. The time range of interest (200 ms – 400 ms) is shaded in yellow. Lower row depicts the MMN wave (blue) calculated by subtracting ERP of match from mismatch, and the confidence interval of 95% estimated from the baseline is depicted with grey line dashed line. The area under the confidence interval within the time range of interest is depicted in grey.

Evoked power from band groups were employed in the decoding algorithm (Figure 6). Decoding performance of band groups were compared in Figure 7, for each condition: Perception and Imagery during task, and Random Perception which is one of the passive control tasks. The decoding performance was significantly different among Conditions (F(2, 315) = 65.65, p = 1.47*10^−24^, 2-way ANOVA) and Band groups (F(4,315)=145.15, p=3.47*10^−70^, 2-way ANOVA) with a significant interaction (F(8,315) = 1.98, p=0.04, 2-way ANOVA) (Figure S5). Decoding performance of Perception was higher than Imagery (p = 1.28 * 10^−14^, Bonferroni) and Random Perception (p = 1.35*10^−23^, Bonferroni), and Imagery was higher than Random Perception (p = 0.02, Bonferroni). For the band groups, decoding performance was higher when two bands were used, theta-alpha, theta-beta, and alpha-beta, than when using lower band group and all bands (Figure S5), indicating additional information from more bands was not helpful to differentiate each pitch. However, an interaction between condition and band groups revealed that the theta-beta model showed the highest performance in Perception and Imagery, but in the Random Perception theta-alpha had the highest performance. The decoding accuracy of the band pairs in the lower frequency showed earlier peak in the performance, and peaking again after the decrement in performance during Perception and Imagery with significant interaction between Condition and Time period only with theta-beta band results (F(4,197) = 2,78, p=0.03, 2-way ANOVA, Figure S6).

**Figure 6.**
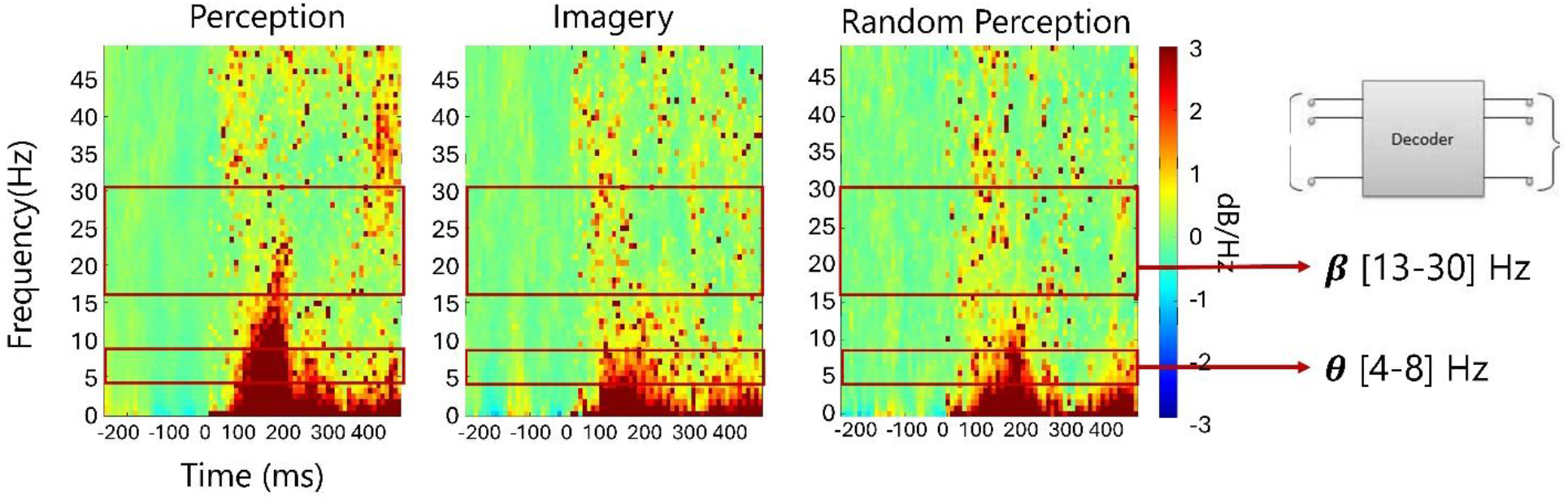
Evoked Power distribution during Perception, Imagery, and Random Perception (example from E3 note from participant 22). The frequency bands corresponding to theta and beta, used as input for the decoder, is indicated as an example.

**Figure 7.**
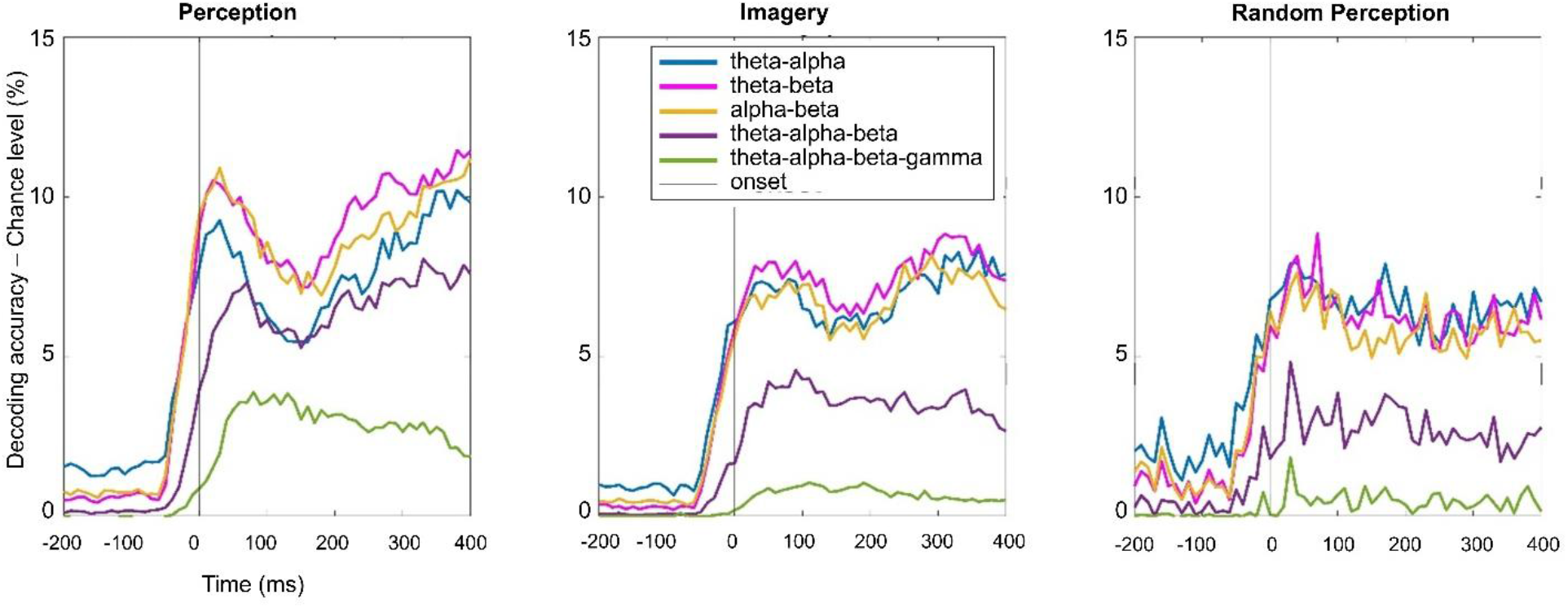
Decoding performance for band pairs in lower band and band combinations. The chance level was estimated by 10 iterations of random permuted model for all band groups. Note that theta-beta model was recomputed to match the configuration to other groups’ model.

Decoding results of the optimal band group are shown in the Figure 8. The decoding results are depicted with decoding accuracy and chance level, with 95% confidence intervals. Overall, the decoding accuracy was higher than the chance level from around 50 ms prior to the onset, resulted from the decoding algorithm using sliding window. The confidence interval between the decoding accuracy and the chance level started to separate from the stimulus onset (time zero on the graph) and maintained that separation for most of the time after the onset. The cluster-based permutation test showed decoding accuracy higher than chance level immediately after the onset (p <0.05).

**Figure 8.**
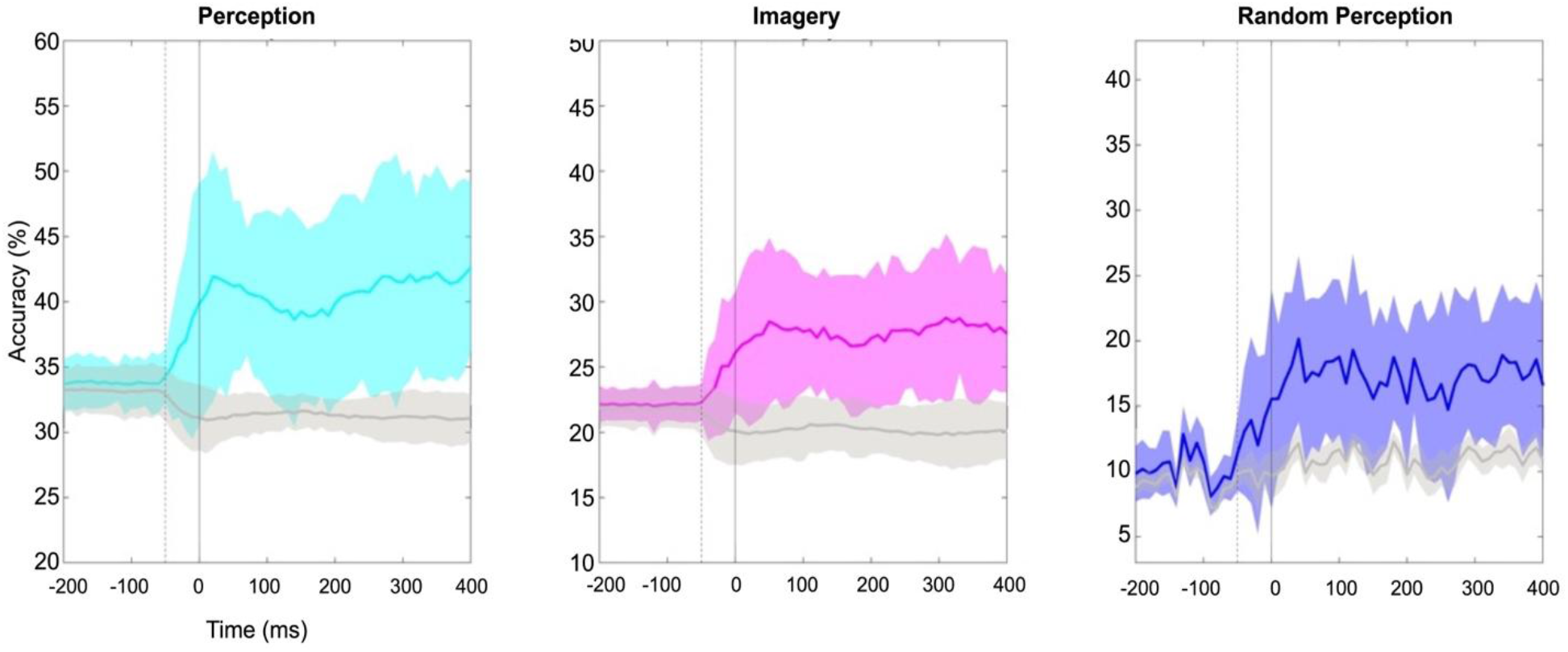
Decoding results for each of the three conditions using theta and beta band power. The decoding accuracy and its 95% confidence interval are depicted with colored lines and shades: cyan for Perception, magenta for Imagery, and blue for Random Rerception, and those of the chance level were depicted with grey color lines and shades. The solid black line is the stimulus onset, but note that the decoding window includes spectral features from −50 ms (dotted line) due to the sliding time window.

The performance was not improved when more EEG channels were used beyond the originally selected temporal channels, namely whole brain, and all channels within each hemisphere tested separately, showing the temporal channel set yielded significantly higher accuracy than other channel combinations (F(3,84) = 98.8 p=1.83*10^−27^, 2-way ANOVA, Figure S7), showing the decodable brain activity for each musical pitch was most informative when coming from near the auditory region related temporal EEG channels from both hemispheres.

## Discussion

The present study examined how specific pitch information is represented during perception and imagery using univariate analyses of EEG evoked responses, and decoding analyses with evoked power across oscillatory frequency bands. A key strength of the experimental design was the adaptation of the familiar-melody task from Herholz et al. (2008), which constrained both the content and timing of each imagery event, allowing this covert process to be examined as a time-locked neural response in EEG. The findings suggest that musical imagery recruits neural processes overlapping with perception, as shown by the parallel evoked potential responses in both perception and imagery, similar MMN responses when the final tone did not match the imagined tone, and correlation across EEG surface topographies. Across all univariate analyses, pitch imagery evoked time-locked neural responses resembling perception, particularly within right temporal electrodes. Finally, the multivariate analysis revealed that pitch-specific information could be decoded from low-frequency oscillatory EEG activity in bilateral temporal regions. Importantly, the present results were observable directly in the electrophysiological signal, rather than by interpreting the inferred encoding or decoding model weights as indices of neural representation, which require additional assumptions and transformations (Haufe et al., 2014).

The similarity in ERP results for Perception and Imagery (Figure 4) is consistent with a broad literature showing that imagery and perception rely on overlapping sensory representations across modalities (Zatorre and Halpern, 2005; Dijkstra et al., 2019; Gu et al., 2019). Beyond spatial overlap, previous studies have shown that neural activity associated with music perception can be reinstated during imagery, in a temporally synchronized manner (Regev et al., 2021; Jakubowski et al., 2019), and that imagery can elicit time-locked auditory components such as N1 and P2 (Janata, 2001; Schaefer et al., 2013; Proverbio et al., 2023). The present findings (Figures 4 and S3) extend this literature by suggesting that such temporal correspondence also applies to individual pitch representation, when imagery timing is experimentally controlled. However, this overlap does not imply equivalence: perception is primarily driven by bottom-up sensory input, whereas imagery depends more strongly on top-down internally generated signals (Dijkstra et al., 2017; Herholz et al., 2012; Zatorre, 2024).

The univariate analyses also revealed a hemispheric asymmetry in both perception and imagery, with the right temporal lobe showing stronger temporal-pattern similarity, higher N1 spatial correlation, and larger MMN responses. This pattern is consistent with established evidence for right-hemisphere specialization in pitch processing. Previous fMRI studies have shown that the right auditory cortex is more sensitive to fine-grained pitch changes, whereas left auditory cortex responds more strongly to coarser pitch variation (Zatorre and Belin, 2001; Hyde et al., 2008). Pitch discrimination thresholds have been shown to correlate with the right auditory cortex activation (Bianchi et al., 2017), and pitch discrimination was shown to be impaired by lesions (Johnsrude et al., 2000) or stimulation-induced disruption (Matsushita et al., 2021) in the right temporal lobe. A similar asymmetry has been reported in musical imagery studies (Halpern et al., 2004; Halpern and Zatorre, 1999). Theoretical accounts attribute this asymmetry to differences in spectral versus temporal resolution, such that the right auditory cortex better encodes fine spectral information while the left has faster temporal integration (Albouy et al. 2020; Zatorre, 2024; Flinker et al., 2019, Okamoto and Kakigi, 2015). The present findings support this specialization beyond externally driven perception to internally generated pitch representations. Greater similarity between imagery and perception in the right hemisphere, together with larger right-hemisphere N1 and MMN responses during imagery, suggests that the right hemisphere may support more vivid or perceptual-like representations of pitch height in the absence of external sound.

The multivariate decoding analyses further showed that low-frequency evoked power carries robust information about individual pitch information in all three Conditions, with accuracy decreasing from Perception to Imagery to Random Perception (Figure 7). This pattern is theoretically informative. The highest decoding performance in Perception likely reflects the combined contribution of sensory input and predictive structure (since the melodies were familiar, each note was strongly predicted by the previous one), whereas lower performance in Random Perception suggests externally presented tones without coherent melodic expectation support weaker pitch-specific coding. That Imagery exceeded Random Perception is particularly notable, indicating that internally generated pitch representations in a familiar musical context may be more structured than neural responses to externally presented but less prediction involving stimuli.

The frequency-band analysis (Figure S5) showed that the frequency-band combinations contributing most to decoding differed across conditions: theta-beta was most informative for Perception and Imagery (Figure 8), whereas theta-alpha was more informative for Random Perception. This pattern suggests that imagery and perception for familiar, predictable sequences share oscillatory mechanism, likely related to predictive processing and the maintenance of pitch information, while Random Perception engages a somewhat different balance of processes. The temporal profile of theta-beta decoding supports this interpretation: in both Perception and Imagery, decoding accuracy emerged early (though after the time-locked stimulus onset), declined during the mid-trial, and increased again later (Figure S6). This profile broadly resembles previous MEG findings showing early pitch-height coding around 100-250 ms, followed by later processing stages associated with higher-order auditory organization such as tonal hierarchy and pitch chroma equivalence (Sankaran et al., 2020; Abrams et al., 2025; Chang et al., 2025). The present findings extend this literature by suggesting temporally structured pitch representations are not limited to heard sounds but can also emerge during internally generated musical events.

The prominence of theta and beta oscillations is also consistent with previous literatures on musical imagery and pitch processing. Theta activity has been linked to auditory-motor interaction, auditory working memory, and the maintenance of internally generated sound representations (Behroozmand et al., 2015; Albouy et al., 2017; Gelding et al., 2019). Beta activity has been associated with top-down prediction, sensorimotor coordination, and the stabilization of internally maintained representations (Arnal and Giraud, 2012; Fujioka et al., 2015). Theta- and beta-band activity in dorsal regions (preSMA, superior parietal cortex, and the dorsolateral prefrontal cortex) has been shown to differentiate pitch perception from pitch labeling in relative-pitch musicians (Leipold et al., 2019). Against this background, our findings suggest that theta-beta dynamics contribute to the retention, prediction, and internal reconstruction of pitch information during both perception and imagery, carrying pitch-specific content rather than reflecting only generic imagery-related activation.

The present findings also contribute to decoding work on musical imagery. Previous studies have mainly focused on higher-order perceptual constructs such as melody identity, melodic contour, scale degree, or tonal organization from the auditory cortex, somatosensory and motor areas (Regev et al. 2021, May et al., 2023; Quiroga-Martinez et al., 2024), or melodic expectations in EEG responses using temporal response function modeling (Marion et al., 2021). By contrast, the present study focused on individual pitch information, a more elementary but perhaps more methodologically difficult feature to isolate during imagery. Our task addresses this fine-grained content by combining familiar melodic continuation with externally paced timing. In this respect, the study extends prior literature by showing that specific pitches can be decoded from low-frequency EEG activity during imagery under relatively controlled conditions.

One caveat is that all melodies were in the third octave, so the decoded pitch information cannot be unambiguously attributed to pitch height vs pitch chroma. The temporal profile is suggestive, earlier decoding aligns with pitch height, and later decoding with pitch chroma (Chang et al., 2025), but disentangling them definitively will require varying octave in future work.

Several limitations should be considered. First, although the visual-control procedure substantially reduced the influence of visually evoked responses (Figures S1 and S2), some residual visual contribution may have affected the evoked responses; however, the visual stimulus could not have affected the MMN response or pitch decoding, since it was identical on every trial. Second, the focus on temporal-lobe EEG channels was theoretically motivated and remained plausible after visual correction (Figure 4), but it excluded other scalp regions, including central channels known to contribute to auditory evoked responses and pitch processing (Näätänen et al., 2007). Third, the relatively low spatial resolution of EEG limits firm anatomical interpretation (Michel and Murray, 2012; Luck, 2014). Relatedly, hemispheric comparisons of oscillatory decoding were constrained by the small number of temporal electrodes per hemisphere, preventing more reliable lateralized decoding. Thus, although we found right-hemisphere dominance in temporal evoked responses, hemispheric asymmetries in oscillatory pitch coding remain to be clarified.

More broadly, the present findings should not be taken to imply that imagery and perception are identical processes. Overlap in evoked responses and decoding patterns can coexist with different network organization or computational demands. For example, musical imagery involves stronger functional connectivity between the anterior right temporal cortex and frontal regions than perception, despite shared activation in secondary auditory cortex (Herholz et al., 2012). In vision, Mechelli et al. (2004) showed perception relies more strongly on occipito-temporal pathways, whereas imagery engages parietal-frontal cortex circuitry, indicating distinct cortical interactions during bottom-up and top-down processing. The present results fit this framework: imagery appears to preserve perceptual-like pitch representations in temporal dynamics and low-frequency oscillations but likely achieves this through partially distinct large-scale network interactions.

## Author Contributions

M.-y.C. designed research with supervision from R.J.Z. and A.R.H; M.-y.C. and C.V.B. performed research; M.-y.C. analyzed data; M.-y.C. wrote the paper; R.J.Z and A.R.H edited and revised the paper. All authors discussed the results and approved the final manuscript.

## Acknowledge

This research was supported by research grants to R.J.Z. from the Canadian Institutes of Health Research and from the Natural Sciences and Engineering Research Council of Canada to R.J.Z, and by the Molson Neuro-Engineering Award from the Montreal Neurological Institute to M.-y.C (2024). R.J.Z. is supported by the Canada Research Chair program and by the Grand Prix Scientifique from the Fondation pour l’Audition (Paris).

