## Supplementary Materials for "Decoding Individual Musical Pitches in Perception and Imagery using Evoked Theta and Beta Power in EEG"

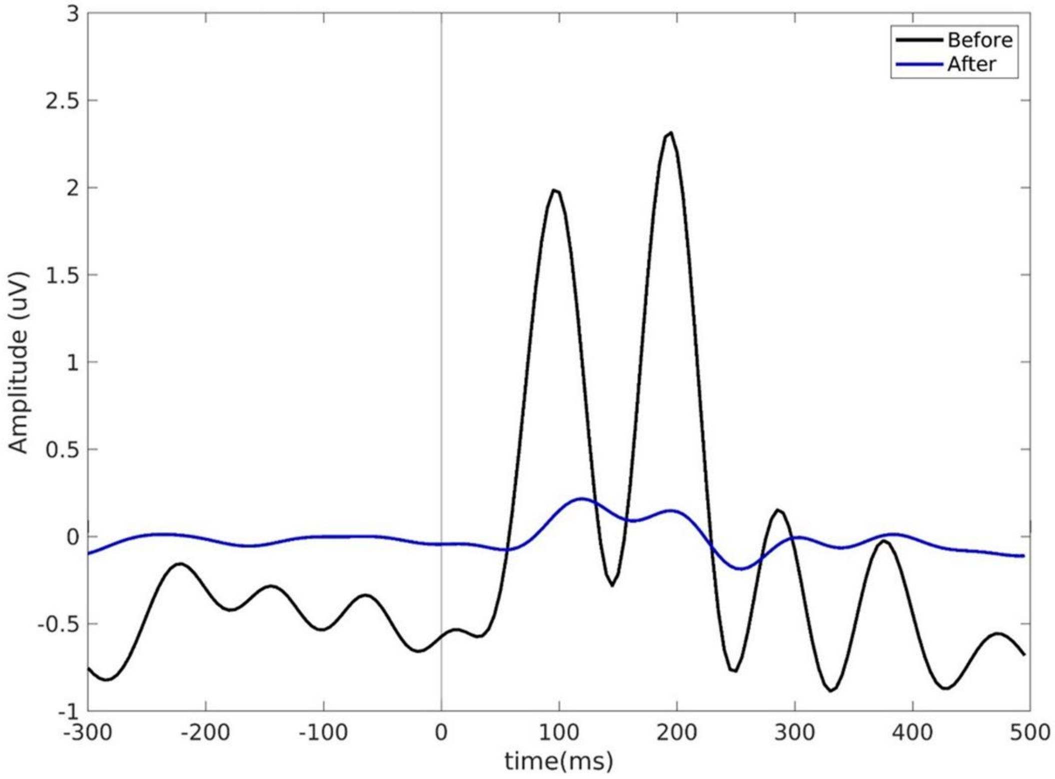


Figure S1. Visual correction before and after the IC correction. ERP plot during visual control session from the occipital channels (Oz, O1, O2, POz, PO3, PO4, PO7, PO8) before (black) and after visual correction (blue), showing the attenuated ERP component from visual stimuli after visual correction.

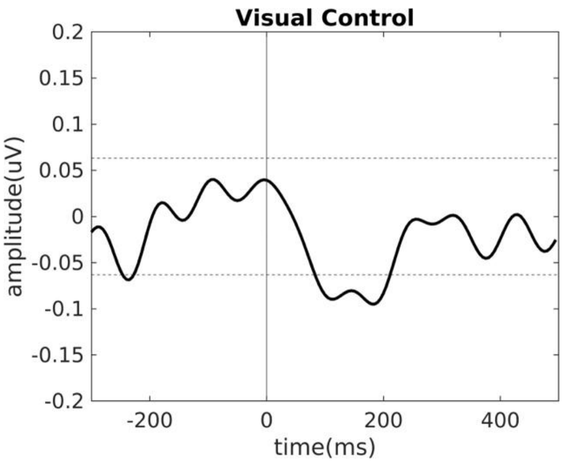


Figure S2. ERP plot during visual control session from the Temporal channels. 95% confidence interval is depicted with the dot line, showing a smaller area of ERP exceeding this line compared to Perception and Imagery.

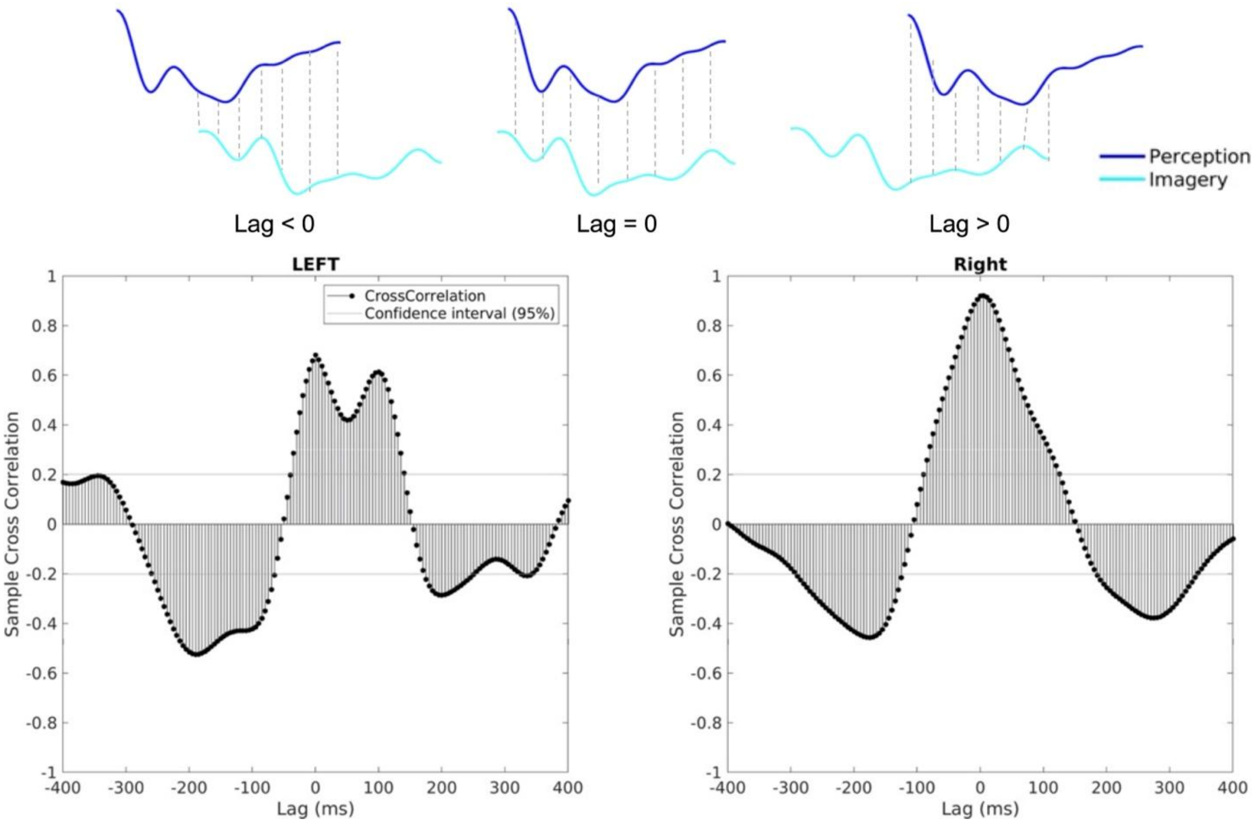

Figure S3. Cross correlation of ERP patterns between Perception and Imagery. Positive lag means the correlation between perception and imagery when the time lag applied to imagery, and vice versa.

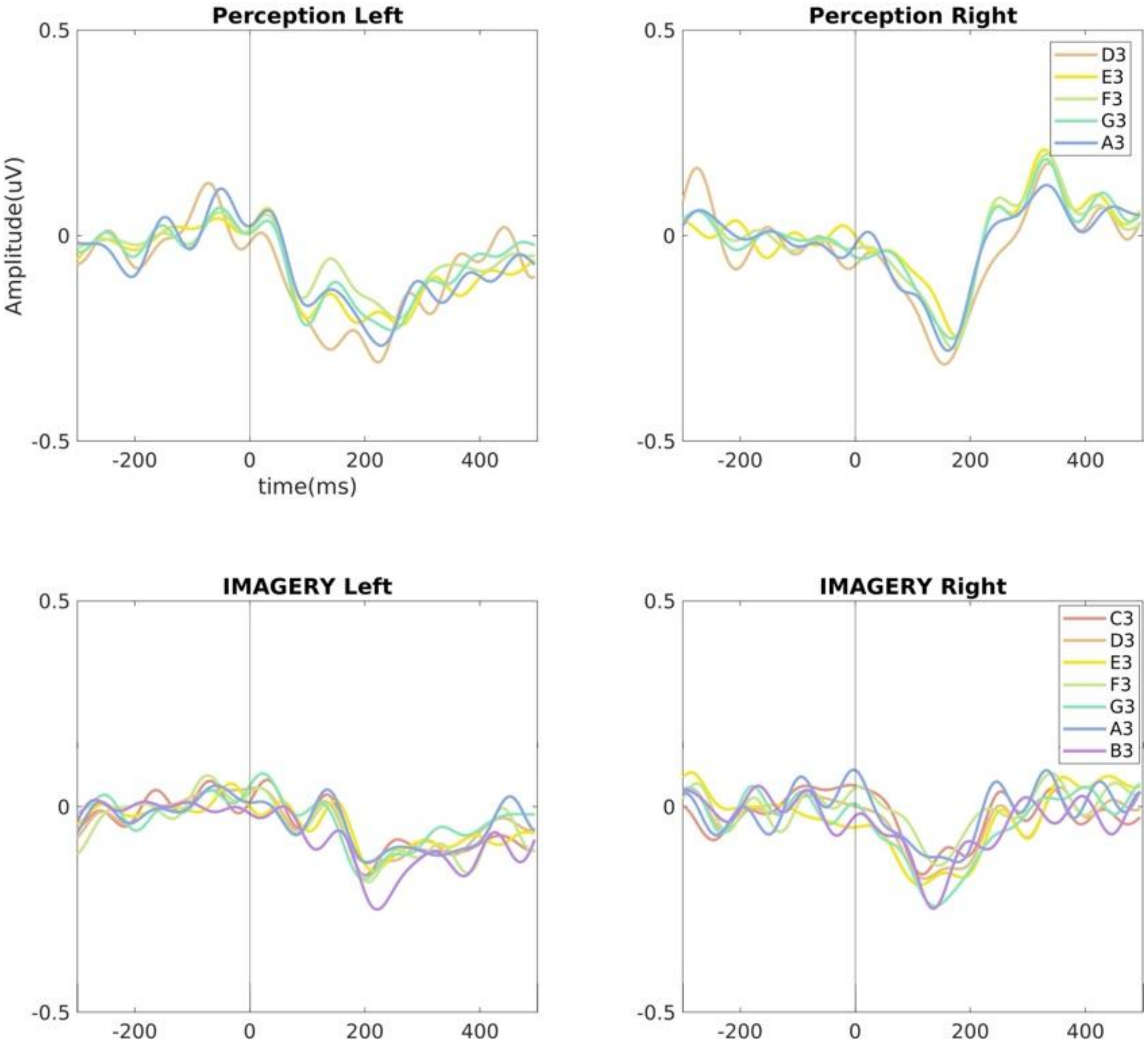


Figure S4. ERPs from Figure4 when including each pitch height.

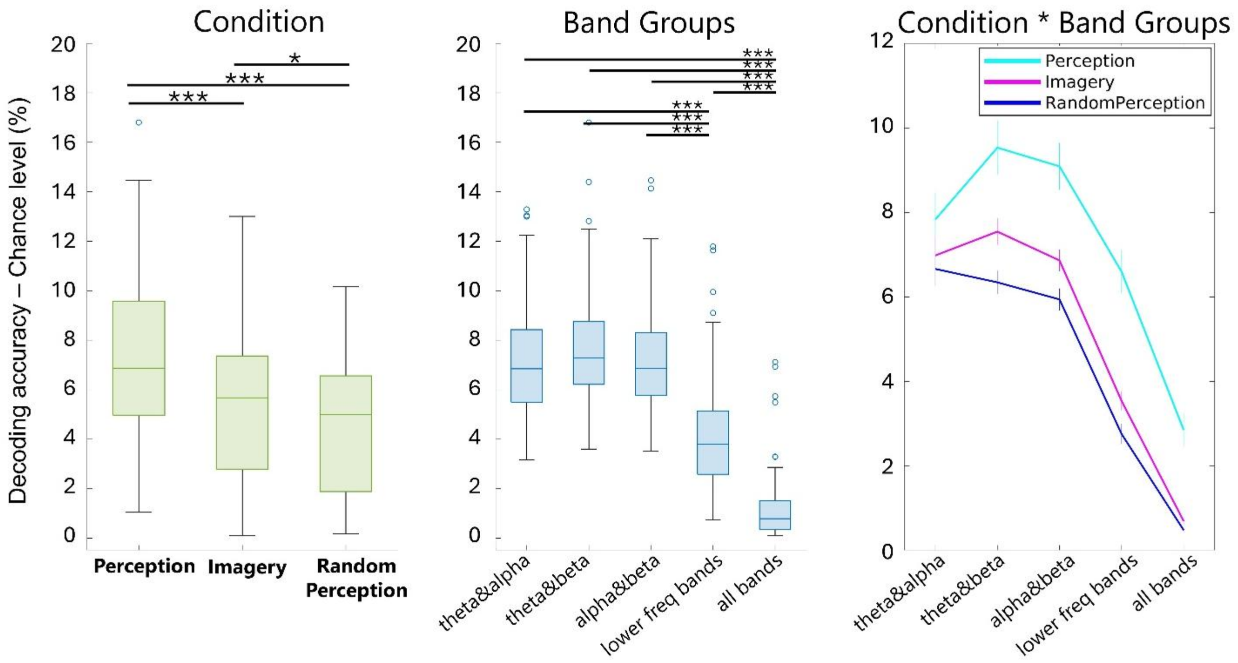


Figure S5. Summary of 2-way ANOVA results for decoding performance of each condition and each band group.

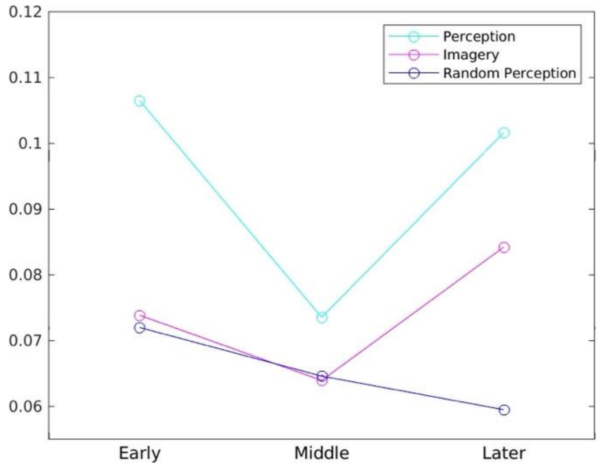


Figure S6. 2-way ANOVA results of theta & beta band decoding performance comparing the interaction between Conditions and Time Period, showing the 2 peak times for Perception and Imagery.

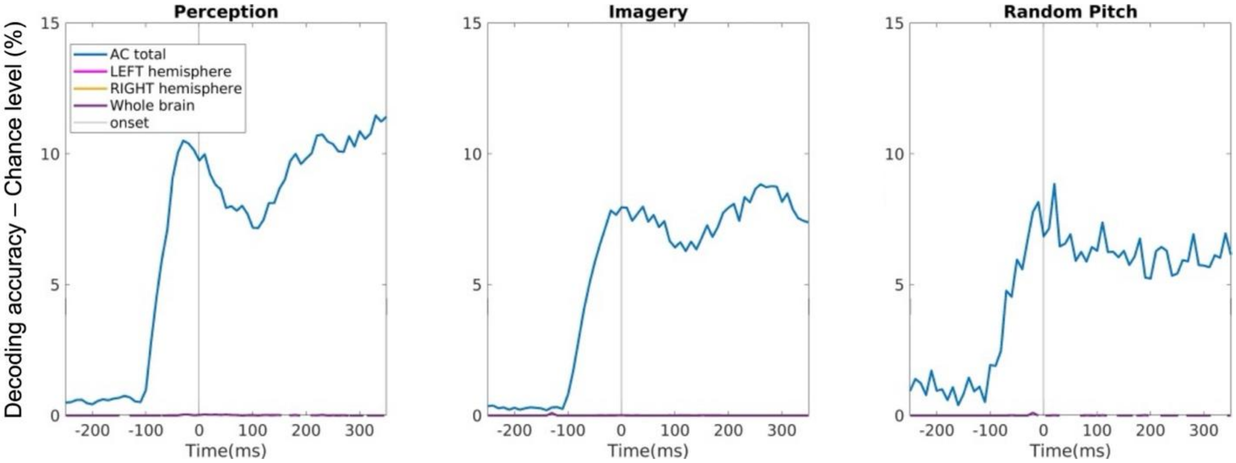


Figure S7. Decoding performance comparison using the theta-beta band pair decoding results from Auditory channels, whole channels from left hemisphere and right hemisphere, and the whole brain EEG channels.
